# EMG1 methyltransferase activity regulates ribosome occupancy at viral uORFs

**DOI:** 10.1101/2024.07.23.604771

**Authors:** Elena M. Harrington, James C. Murphy, Julie L. Aspden, Adrian Whitehouse

**Affiliations:** School of Molecular and Cellular Biology, Faculty of Biological Sciences, University of Leeds, Leeds, LS2 9JT, United Kingdom; Astbury Centre for Structural Molecular Biology, University of Leeds, Leeds, LS2 9JT, United Kingdom; LeedsOmics, University of Leeds, Leeds, LS2 9JT, United Kingdom

**Keywords:** KSHV, EMG1, uORFs, specialised ribosome

## Abstract

Viruses do not encode their own translational machinery and rely exclusively on the translational machinery of the host cell for the synthesis of viral proteins. Viruses have evolved diverse mechanisms to redirect the cellular translation apparatus to favour viral transcripts. A unique mechanism employed by Kaposi’s sarcoma-associated herpesvirus (KSHV) involves manipulation of cellular ribosome composition, producing virus-induced specialised ribosomes. These ribosomes scan through KSHV uORFs in late lytic genes, allowing efficient translation of downstream main KSHV ORFs. Herein, we highlight the enhanced association of the ribosomal biogenesis factor EMG1 with precursor-40S ribosome complexes during KSHV lytic replication. Depletion of EMG1 results in significantly reduced expression of viral proteins and progression through the lytic cascade, culminating in a dramatic reduction of infectious virus production. We further demonstrate that the methyltransferase activity of EMG1 is required for effective regulation of translation of KSHV uORFs in late lytic genes.

## Introduction

Protein synthesis is a highly dynamic process, constantly responding to changes in the environment to maintain cell homeostasis^1,2^. Historically, changes in protein synthesis were thought to be due to either gene regulation or altered characteristics of mRNAs whereas ribosomes were considered as ubiquitous, static, machine-like entities with a passive role in translation^3^. However, emerging evidence supports an increasingly complex and dynamic network of translational regulation. Ribosome specialisation offers the potential to selectively translate specific subsets of mRNAs or ORFs leading to altered levels of protein expression and distinct phenotypes across cell and tissue types^3,4^. Mechanisms leading to heterogenous populations of ribosomes within the cell can involve changes in ribosomal protein stoichiometry^5^, paralogue switching^6^, posttranslational modifications^7^, addition of ribosome-associated proteins^8,9^ and changes in rRNA genetic variation and modifications^10^. Evidence of specialised ribosomes is now being uncovered in differing tissue types, developmental stages, or due to environmental factors such as stress, temperature and pathogens^3^. This is further highlighted in ribosomopathies, a group of diseases caused by mutations or loss of ribosomal biogenesis proteins that lead to reduced or altered ribosome functioning^11^.

The rapidly developing concept of specialised ribosomes poses an intriguing question as to whether viruses co-opt this cellular mechanism to enhance translation of their own mRNAs. Viruses lack their own translational machinery so instead rely on host ribosomes to synthesise their proteins, competing with endogenous host transcripts. Therefore, viruses have evolved diverse mechanisms to ensure translational efficiency of viral mRNA above and beyond that of cellular mRNA^12,13^. These processes serve to redirect the translation apparatus to favour viral transcripts, often coming at the expense of the host cell.

Kaposi’s sarcoma-associated herpesvirus (KSHV) is a large double stranded DNA virus associated with Kaposi’s sarcoma (KS) and two lymphoproliferative disorders; multicentric Castlemans disease (MCD) and primary effusion lymphoma (PEL)^14,15^. KSHV, like all herpesviruses exhibits a biphasic lifecycle, encompassing latent persistence and lytic replication phases^16^. During latency the virus remains transcriptionally dormant, with expression limited to a few latent genes, enabling long term persistence of the viral episome^17^. Under various stimuli, the KSHV episome can be reactivated into lytic replication, leading to the ordered expression of more than 80 viral proteins, culminating in the production of infectious virions^18^. This dramatic shift in viral gene expression during lytic replication from the relative dormancy of latency puts a high demand on the cellular translational machinery and one way KSHV circumvents this issue is by producing virus-specific specialised ribosomes which preferentially translate viral mRNAs^19^. Quantitative proteomic affinity pulldowns have previously identified changes in the stoichiometry and composition of pre-40S ribosomal complexes between latent and lytic KSHV replication cycles. Analysis identified two ribosomal complexes that have increased association with the pre-40S ribosomal subunit during lytic KSHV replication, namely the BUD23-TRMT112 and NOC4L-NOP14-EMG1 complexes. In addition, ribosome profiling showed that these virus-induced specialised ribosomes reduce the translation of specific uORFs present in KSHV late lytic transcripts, which in turn enhances the translation of the downstream CDSs^19^. Interestingly, both the BUD23-TRMT112 and NOC4L-NOP14-EMG1 complexes contain methyltransferases, which methylate specific residues on the 18S rRNA. BUD23 mediates the N7-methylation of G1639^20^, whereas EMG1 catalyses the N1-methylation of 1248U^21^. Notably, both BUD23 and EMG1 also have additional roles in ribosome biogenesis acting as an assembly scaffold proteins^22,23^.

Herein we aimed to examine the specific role of EMG1 in virus-induced specialised ribosome function. We show enhanced association of EMG1 with the pre-40S ribosomal subunit during KSHV lytic replication and confirm EMG1 is essential for KSHV lytic replication and infectious virion production. Importantly, we demonstrate that the methyltransferase activity of EMG1 is necessary for reduced occupancy at viral uORFs allowing enhanced translation of the downstream viral CDS. This provides evidence that modification of rRNA can affect the activity of ribosomes during infection.

## Results

### EMG1 has increased association with the pre-40S ribosomal subunit during KSHV lytic replication

Quantitative proteomic pulldown studies have previously characterised changes in pre-40S ribosomal complexes during KSHV infection^19^. The NOC4L-NOP14-EMG1 complex was shown to have substantially increased stoichiometry with pre-40S complexes during KSHV lytic compared to latent replication cycles. EMG1 is the methyltransferase within the complex which catalyses the N1-methylation of 1248U residue on the 18S rRNA^22^, therefore we aimed to determine whether the catalytic activity of EMG1 is required for KSHV-mediated specialised ribosome function.

To firstly confirm an increased association between EMG1 and the pre-40S ribosomal subunit during KSHV lytic replication, FLAG-TRAP affinity pulldowns were performed using a TREx BCBL1-Rta cell line stably expressing FLAG-2xStrep-PNO1, during latent and lytic replication phases. Western blot analysis confirmed a 3-fold increased association between EMG1 and PNO1 during KSHV lytic replication, compared with latently-infected cells, whilst no change in association was observed in the control core ribosomal protein, eS19, confirming the previous quantitative proteomic results (**Fig. 1a**). KSHV ORF11 has been previously identified as the key viral mediator in the production of KSHV specialised ribosomes^19^. Therefore, FLAG-TRAP affinity pulldowns were performed using a TREx BCBL1-Rta cell line stably expressing FLAG-ORF11 during both latent and lytic replication phases to assess whether KSHV ORF11 interacts with the EMG1-NOC4L-NOP14 complex. Western blot analysis confirmed that the EMG1-NOC4L-NOP14 complex associates with ORF11, aligning with the hypothesis that ORF11 is recruiting these proteins to the pre-40S ribosomal subunit during the early stages of lytic replication. Moreover, the interaction between FLAG-ORF11 and the EMG1-NOC4L-NOP14 complex during latency suggests no other KSHV lytic proteins are required for this association to occur (**Fig. 1b**).

**Figure 1.**
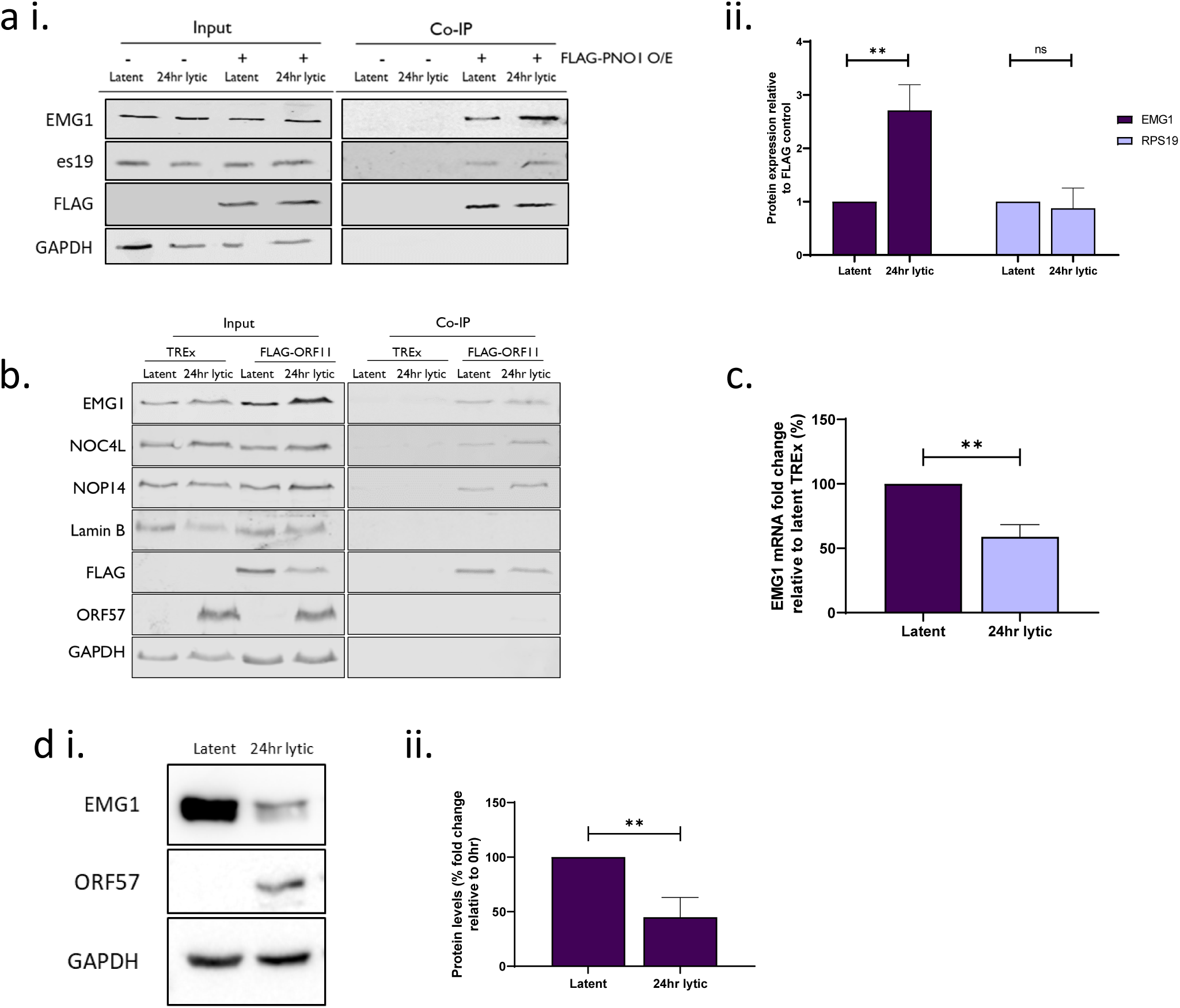
Increased association of EMG1 with pre-40S complexes during KSHV lytic replication. TREx BCBL-Rta cells stably expressing a FLAG-PNO1 overexpression construct, or wildtype TREx BCBL-Rta cells, remained latent or were induced for KSHV lytic replication for 24 hours. **ai)** FLAG-TRAP elutions were performed on the whole cell lysates and probed via western blot alongside the whole cell lysate inputs with EMG1, FLAG, eS19 and GAPDH-specific antibodies (n=3). **ii.** Densitometric analysis of the FLAG-Trap pulldowns, normalised to the levels of the bait protein PNO1 (FLAG) (n=3). TREx BCBL-Rta cells stably expressing a FLAG-ORF11 overexpression construct, or wildtype TREx BCBL-Rta cells remained latent or were induced for KSHV lytic replication for 24 hours. **b)** FLAG-TRAP elutions were performed on the whole cell lysates and probed via western blot alongside the whole cell lysate inputs with EMG1, NOC4L, NOP14, Lamin B, FLAG, ORF57 and GAPDH-specific antibodies (n=3). TREx BCBL-Rta cells remained latent or were reactivated for 24hr. **(c)** RNA and **(di)** protein extracted from cell lysates probed for EMG1 expression, normalised to GAPDH (n=3). **dii)** Densitometric analysis of EMG1 protein expression, normalised to GAPDH (n=3). Data presented as mean ± SD. Asterisks denote a significant difference between the specified groups (\**p* ≤ 0.05, \*\**p* < 0.01 and \*\*\**p* < 0.001).

To examine whether the increase association of EMG1 was due to an upregulation of expression during infection, EMG1 mRNA and protein levels were assessed at 24 hours post reactivation. Surprisingly, whilst EMG1 has an increased association with the pre-40S during KSHV lytic replication, the overall mRNA and protein levels of EMG1 are reduced during lytic replication (**Fig. 1c–d**), suggesting that the increased association of EMG1 is directly driven by the virus at early stages of replication, circumventing KSHV SOX-mediated host cell shutoff of the EMG1 mRNA. These results suggest that KSHV, via the ORF11 protein, enhances the association between EMG1 and pre-40S during ribosomal biogenesis in the early stages of lytic replication.

### EMG1 is essential for efficient KSHV lytic replication

To determine whether the increased association of EMG1 with the pre-40S subunit is important for KSHV lytic replication, TREx-BCBL1-Rta cells were stably transduced with lentivirus-based EMG1 targeting shRNAs, depleting EMG1 mRNA levels by 70% (KD1) and 60% (KD2) and protein levels by 98% (KD1) and 96% (KD2), respectively (**Fig. 2a-b**). To confirm that depletion of a host ribosome biogenesis factor had limited effect on TREx BCBL1-Rta, a series of assays measuring protein turnover, ribosome population and translational capacity were performed comparing the two EMG1 depleted TREx-BCBL1-Rta cell lines and a scrambled control. Global protein turnover was measured using CDK1 as a marker, with western blot analysis showing no significant change in CDK1 protein levels in scrambled shRNA treated control cells compared to cells depleted of EMG1 (**Fig. 2c**). Polysome profiling was also used to compare the ribosome population, with results showing that levels of 80S ribosomes and polysomes upon EMG1 knockdown are similar to control, indicating that loss of EMG1 had little effect on ribosome biogenesis or overall ribosome levels in KSHV-latently infected cells (**Fig. 2d**). Finally, EMG1 depletion had no significant effect on global translational output, as assayed by labelling nascent proteins with click-iT® L-Azidohomoalanine and quantifying by flow cytometry (**Fig. 2e**). Based on these assays, no significant adverse effects to TREx-BCBL1-Rta ribosome population and translational capacity was observed upon EMG1 depletion.

**Figure 2.**
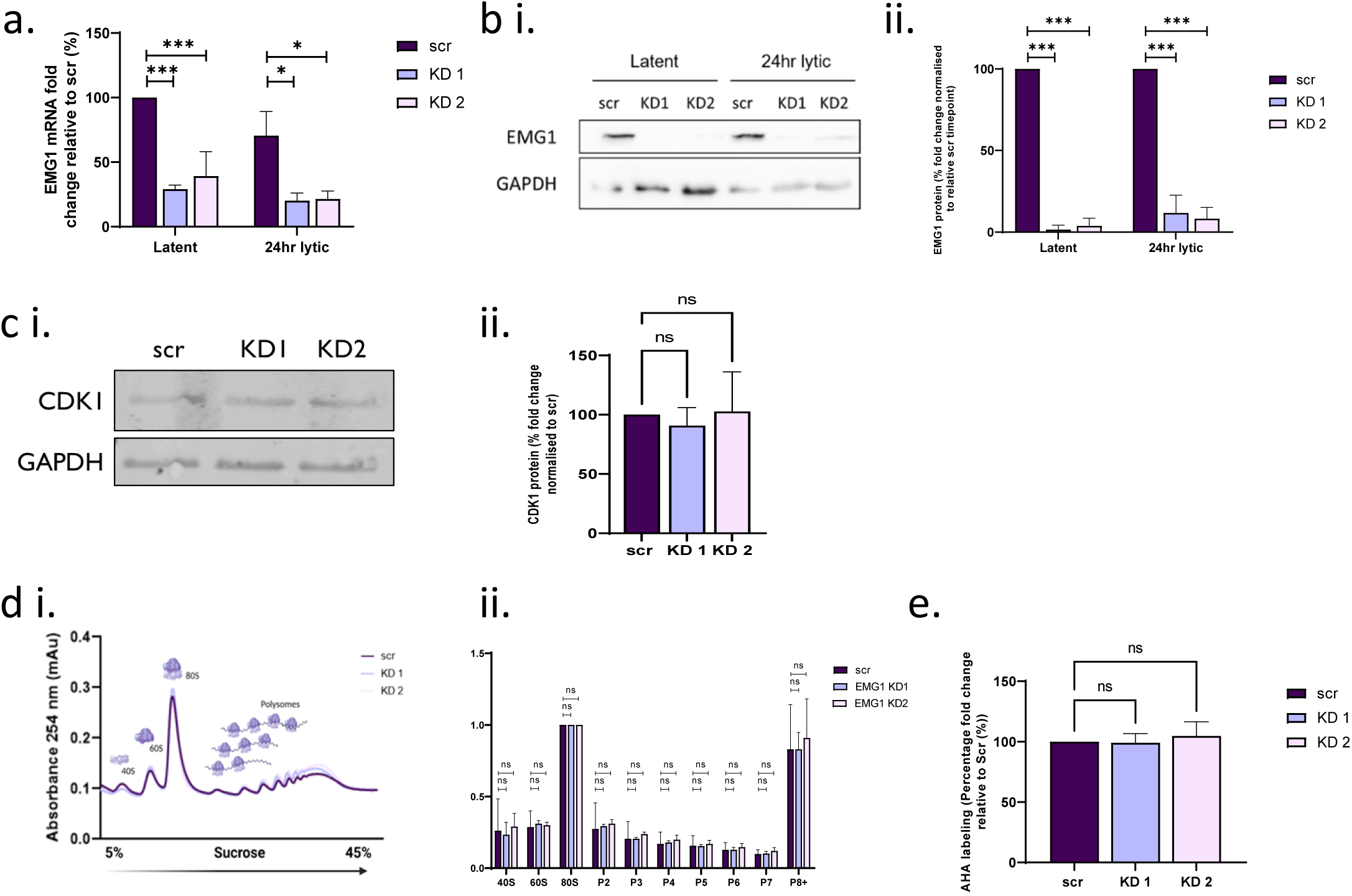
EMG1 depletion does not affect global translation in KSHV-infected TREx BCBL-Rta cells. TREx BCBL-Rta cells containing shRNAs targeting EMG1, or a scr shRNA, were used to study the effects of EMG1 depletion. mRNA **(a)** and protein **(bi)** expression of EMG1 in EMG1 depleted TREx-BCBL-Rta cells compared to the scr control during latent and lytic KSHV replication (n=3). **bii)** Densitometric analysis of EMG1 protein expression in TREx BCBL-Rta cells depleted of EMG1 compared to scr control, normalised to GAPDH (n=3). **ci)** Protein lysates of latent TREx BCBL-Rta cells depleted of EMG1 or containing a scr control plasmid were probed via western blot with CDK1 and GAPDH antibodies (n=3). **cii).** Densitometric analysis of CDK1 expression in EMG1 depleted cells, normalised to GAPDH (n=3). **di)** Polysome profiling traces of latent TREx BCBL-Rta cells depleted of EMG1 or containing scr control plasmid (n=3). **dii).** Densitometric analysis of each polysome fraction relative to the 80S subunit. The densitometric value of each peak was measured in triplicate and averaged on each biological repeat (n=3). **e)** A methionine chase experiment was used to measure global translation by depleting methionine from the cells, then labelling nascent peptides with click-iT L-Azidohomoalanine (AHA). Abundance of AHA was measured via flow cytometry using an alkyne Alexa Fluor 488 (n=3). Data are presented as mean ± SD. Asterisks denote a significant difference between the specified groups (\**p* ≤ 0.05, \*\**p* < 0.01 and \*\*\**p* < 0.001).

To assess what effect preventing the enhanced association of EMG1 with pre-40S ribosomal complexes had upon KSHV lytic replication, the transcription and translation of viral early and late genes was assessed in EMG1-depleted TREx-BCBL1-Rta cell lines compared to a scrambled control. It would be predicted that modification of the host cell ribosome population to produce virus-induced specialised ribosomes during infection, would impact late viral protein production rather than the early stages post reactivation. To this end, mRNA and protein levels of the early KSHV ORF57 and late ORF65 genes were compared at 24 and 48 hour post reactivation. Results showed no statistically significant changes in ORF57 mRNA levels, however a slight increase in ORF65 mRNA was observed during the time course between the scrambled shRNA control and EMG1-knockdown cells (**Fig. 3a**). In contrast, while ORF57 protein levels remained relatively stable 24 hours post lytic induction (**Fig. 3b**), EMG1 depletion significantly inhibited ORF65 protein production, resulting in a 75-76% reduction 48 hours post lytic induction (**Fig. 3c**). RT-qPCR of polysome fractions was then used to observe changes in active translation of viral ORF57 and ORF65 transcripts upon EMG1 depletion compared to scrambled control during KSHV lytic replication. Highly active translating mRNAs, with six or more ribosomes bound, were collected and the levels of ORF57 and ORF65 transcripts in these fractions were quantified. EMG1 depletion resulted in a 35% drop in the level of ORF57 mRNA associated with large polysomes translation after 24 hrs post reactivation, whereas ORF65 was dramatically reduced by 81% (**Fig. 3d**). This suggests EMG1 knockdown results in a reduction in translation of ORF57 and ORF65.

**Figure 3.**
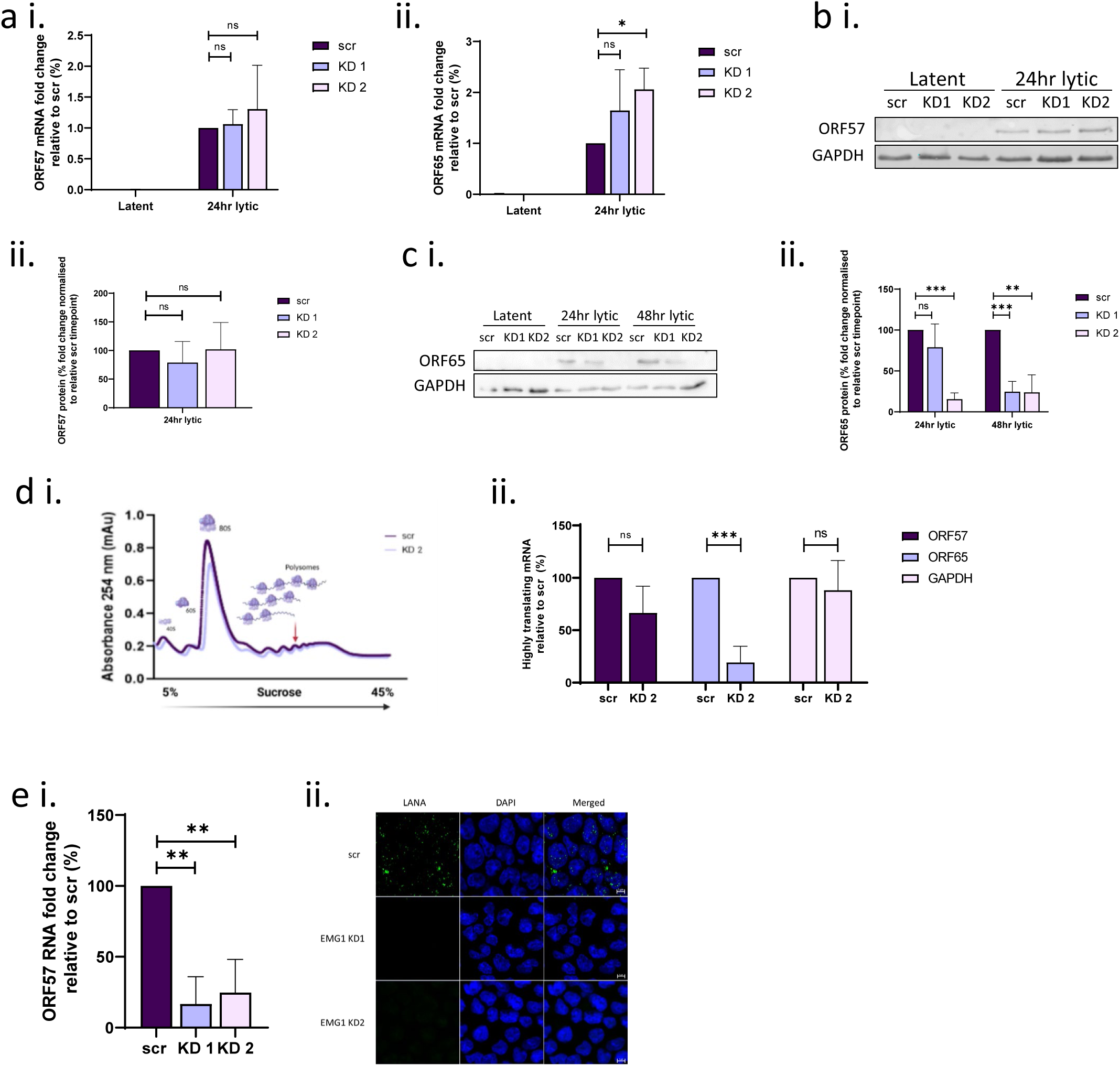
EMG1 Is required for efficient KSHV lytic replication. Latent and 24hr reactivated cell lysates from TREX BCBL-Rta cells depleted of EMG1 or containing a scr control plasmid were analysed for mRNA expression of **(ai)** ORF57 and **(aii)** ORF65 were analysed in latent and 24hr reactivated cell lysates, normalised to GAPDH (n=3). Protein expression of **(bi)** ORF57 and **(ci)** ORF65 were analysed in latent and 24hr reactivated cell lysates (n=3). Densitometric analysis of **(bii)** ORF57 and **(cii)** ORF65 protein expression normalised to GAPDH (n=3). **di)** Polysome profile trace of TREx-BCBL Rta cells depleted of EMG1 or containing a control scr plasmid after 24hr KSHV lytic replication. **dii)** Total RNA was isolated from traces containing five or more polysomes (denoted by arrow) and expression of viral genes ORF57 and ORF65 were analysed via qPCR and normalised to reference genes (n=3). **ei)** Virus released from the TREx-BCBL Rta cell lines was collected and introduced to naïve 293T cells for 48hrs to measure viral reinfection capability. Total RNA was isolated and ORF57 expression was analysed via qPCR, normalised to GAPDH (n=3). **(eii)** Reinfected 293T cells were permeabilised and stained for LANA (green) and DAPI (blue). The cells were mounted on coverslips and visualised using an inverted LSM800 confocal microscope. Data are presented as mean ± SD. Asterisks denote a significant difference between the specified groups (\**p* ≤ 0.05, \*\**p* < 0.01 and \*\*\**p* < 0.001).

Finally, to confirm that EMG1 depletion affected later stages of KSHV lytic replication, infectious virion production was also assessed. Here supernatants of reactivated scrambled and EMG1-depleted TREx-BCBL1-RTA cells were used to re-infect naïve HEK-293T cells and infectious virion production was quantified by qPCR or LANA immunostaining. Naïve cells infected with supernatant from reactivated EMG1-depleted cells resulted in a dramatic loss of infectious virions compared to the scramble control, quantified by qPCR as a reduction of 75-84% (**Fig 3e**). This was further demonstrated by the almost total loss of staining for the viral marker LANA in naive HEK-293T cells, which were re-infected with virus produced from cells depleted of EMG1 (**Fig 3e**). Taken together, these results suggest that EMG1 depletion significantly reduces the translation of late viral proteins which impacts downstream production of new infectious virions.

### EMG1 methyltransferase activity affects ribosome occupancy at viral uORFs impacting translation of the downstream CDS

Ribosome profiling has previously revealed that KSHV specialised ribosomes scan through viral late gene uORFs, overcoming uORF-mediated repression and enhancing translation of the downstream viral main ORFs^19^. To determine whether EMG1, and more specifically its methyltransferase activity, regulate ribosome occupancy at viral uORFs, dual luciferase reporter constructs were utilised, comprising ORF30 and ORF34 5’-UTRs containing uORFs, previously shown to be regulated by KSHV-induced specialised ribosomes^19^. These 5’-UTRs-luciferase reporter constructs were transfected into 293T cells containing a scr shRNA or shRNAs targeting EMG1. Similarly, to TREx-BCBL1-RTA cells, shRNA-mediated depletion resulted in efficient EMG1 knockdown by 80% and 60% (**Fig. 4a**). Notably, EMG1 depletion significantly reduced the expression of the downstream luciferase reporter by 58% and 54% in the presence of the ORF30 5’-UTR containing uORF30, and 63% and 39% in the ORF34 5’-UTR containing uORF34 compared to the scrambled control (**Fig. 4b**).

**Figure 4.**
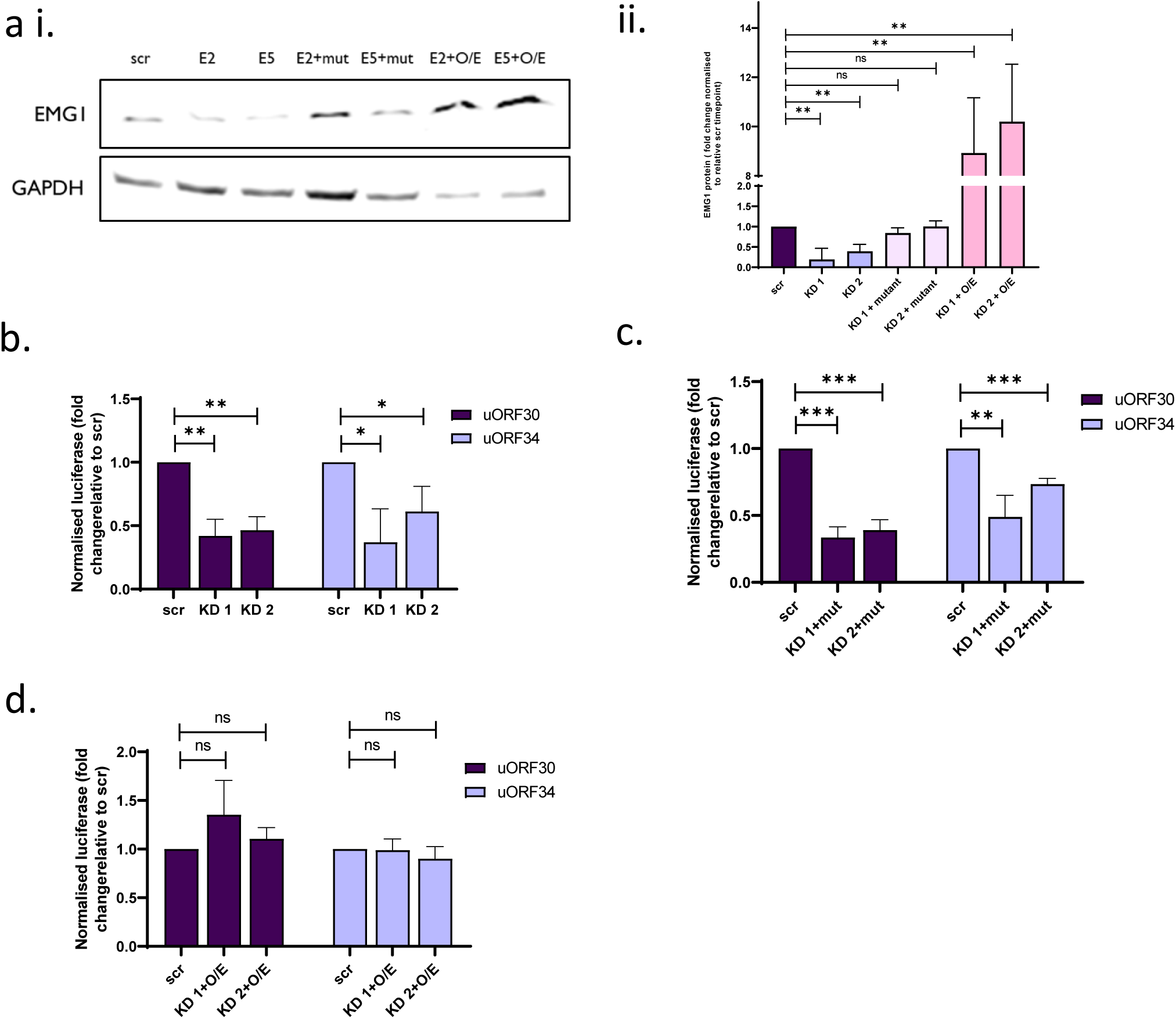
EMG1 methyltransferase activity is required to regulate viral uORF ribosome occupancy. **ai)** Protein lysates of HEK293T cells expressing control shRNA, EMG1 shRNA, catalytically inactive EMG1 (mut) or wildtype EMG1 overexpression plasmids were probed for expression of EMG1 via western blot (n=3). **aii)** Densitometric analysis of EMG1 expression relative to GAPDH in varying HEK293T cell lines (n=3). Dual luciferase reporter constructs containing 5’ untranslated regions (UTRs) were transfected into **(b)** EMG1 depleted cells, **(c)** EMG1 depleted cells rescued with a catalytically inactive EMG1 construct (mut) **(d)** EMG1 depleted cells recovered with a wild type EMG1 overexpression construct (O/E). Luciferase activity was normalised to the scr shRNA control cell line (n=3). Data are presented as mean ± SD. Asterisks denote a significant difference between the specified groups (\**p* ≤ 0.05, \*\**p* < 0.01 and \*\*\**p* < 0.001).

To determine whether the reduction of luciferase activity was dependent on the methyltransferase activity of EMG1, an EMG1 mutant construct was generated, which replaced the previously characterised catalytic loop (RPDI, residue 84-88,) with alanine residues to produce a catalytically inactive EMG1 construct^24^. Cell lines were then generated expressing shRNAs targeting EMG1 and a rescue shRNA resistant construct, either expressing the mutant EMG1 construct or a wild type EMG1 construct. The uORF dual luciferase plasmids were then transfected into each cell line to assess translation regulation. Results showed that rescue of EMG1 knockdown with wild-type EMG1 alleviated this repression. In contrast, expression of the EMG1 catalytic mutant was unable to rescue the reduced expression in the presence of the KSHV 5’-UTRs containing uORFs (**Fig. 4c and d**). Taken together, these findings strongly support the hypothesis that the methyltransferase activity of EMG1 enables reduced translation at viral uORFs allowing enhanced translation of downstream CDSs.

## Discussion

Like all viruses, KSHV lacks it’s own translational machinery and therefore must manipulate the host cell translational apparatus to enable preferential translation of viral mRNAs over host mRNAs^12,19^. Recent studies suggest KSHV produces virus-induced specialised ribosomes by enhancing the recruitment of specific ribosomal biogenesis factors to the pre-40S ribosomal subunit during early stages of lytic infection through the activity of the KSHV ORF11 protein^12,19^. However, the precise role these ribosome-associated factors play is yet to be determined. A clue may lie in the complexes which have been identified using quantitative proteomic pulldown analysis, which shows an an increased association of two complexes during KSHV lytic replication, BUD23 and its co-factor TRMT112, and the NOC4L-NOP14-EMG1 complex. Interestingly, both these complexes contain methyltransferase activities, which modify the 18S rRNA. Specifically, BUD23 methylates m^7^-G1639, whereas EMG1 methylates N^1^-U1248^20,21^. However, it is unknown whether the methytransferase activity of these complexes is essential for KSHV specialised ribosome functioning. Herein, we demonstrate that the methyltransferase activity of EMG1 is essential to regulate KSHV-mediated specialised ribosomes association with viral uORFs which regulate translation of the downstream coding region.

Surprisingly, EMG1 depletion was dispensable for host cell proliferation and had little effect on ribosome biogenesis or global translation in latently KSHV-infected TREx-BCBL1-RTA cells. This may suggest tissue- or cell-specific differences in the requirement of EMG1, similar to other ribosomal biogenesis factors. This is emphasized by a single amino acid mutation (aspartate to glycine) at position 86 within EMG1, which is associated with Bowen-Conradi syndrome^25^. This rare ribosomopathy is characterised by moderate to severe prenatal and postnatal growth retardation, microcephaly, a distinctive facial appearance, profound psychomotor delay, hip and knee contractures and rockerbottom feet^26^. Mutation at residue 86 does not affect it methyltransferase catalytic activity but is thought to lead to its nuclear aggregation and degradation, resulting in the reduced nucleolar recruitment^27^. This emphasises the scaffolding function of EMG1 and presence within the pre-ribosomal complex is essential, whereas the *N*1 – methylation of U1248 on the 18S rRNA is a non-essential modification. This leads to the hypothesis that the methyltransferase activity of EMG1 could be a preferential modification triggered under different conditions, serving to modify the 18S rRNA and contribute to the regulation of translation.

In contrast, here we have shown that EMG1 is essential for efficient KSHV lytic replication and infectious virion production. Notably, EMG1 depletion had little effect on KSHV ORF57 and 65 mRNA levels, however translation and protein production, particularly of the late viral ORF65 protein was dramatically reduced. Taken together, these results demonstrate that EMG1 is essential for enhancing the translation of late viral proteins, resulting in the collapse of the late lytic gene cascade culminating in dramatically reduced levels of infectious virions. This phenotype is similar to that of BUD23, therefore it is likely that both these methytransferases are making modifications to the 18S that result in a cumulative effect that drastically changes which ORFs these ribosomes preferentially translate.

Through ribosome profiling, we have previously shown that KSHV-induced specialised ribosomes prevented occupancy and thereby reduced translation of viral uORFs in late lytic KSHV mRNAs and consequently the efficient translation of the downstream CDS^19^. Here we have investigated whether EMG1 and specifically its methyltransferase activity is involved in this regulatory mechanism. Importantly, luciferase reporter assays confirmed that depletion of EMG1 resulted in a decrease in downstream CDS translation, likely due to an increase in uORF translation. Of key importance here is the finding that the methyltransferase activity of EMG1 is the essential aspect for KSHV replication and infectious virion production. Notably, the 18S rRNA base 1248 has a three-part temporal modification, in which snr33 initially pseudouridinylates the adenosine to a uracil, followed by EMG1 N^1^-methylation and finally Tcr adding an acp3 modification. The modified U1248 base sits in the P site, adjacent to the codon: anti-codon interface, which suggests these modifications are essential for allowing cognate codon:anti-codon pairings^28^. Recent dynamic simulations have suggested that addition of this methyl group stabilises interaction of the ribosome with tRNA^29^. Therefore, without this methylation there may be reduced restriction on codon:anti-codon pairing, allowing more non-canonical base pairing to occur and leaky’ codon pairings passing through the ribosome.

In summary, we show that KSHV enhances the recruitment of EMG1 to newly synthesised pre-40S ribosomal subunits during early stages of KSHV lytic replication. This enables the production of a virally-induced specialised ribosome, which preferentially translates viral late proteins. Mechanistic studies demonstrate that the methyltransferase activity of EMG1 is essential for reduced translation at viral uORFs allowing enhanced translation of the downstream viral CDS.

## Acknowledgements

We thank Professor Jae Jung, University of Southern California School of Medicine, Los Angeles. This work was supported by University of Leeds Mary & Alice Smith Endowed Research Scholarship to EMH and AW, Wellcome Trust to JCM and AW (203826/Z/16/Z), BBSRC to AW and JLA (BB/N014405/1 and BB/X003086/1).

## Author Contributions

Conceptualization (EMH, AW); Data curation (EMH, JCM); Formal Analysis (EMH, JCM, JLA, AW); Funding acquisition (EMH, JCM, JLA, AW); Investigation (EMH, JCM); Writing – original draft (EMH, AW); Writing – review & editing (All authors).

## Methods

### Cell culture

TREx-BCBL1-RTA cells, a B cell lymphoma cell line latently infected with KSHV engineered to contain a doxycycline inducible myc-Rta were a gift from Professor JU Jung (University of Southern California). TREx-BCBL1-RTA cells were cultured in RPMI1640 with glutamine (Gibco), supplemented with 10% Foetal Bovine Serum (FBS) (Gibco), 1% P/S (Gibco) and 100 µg/mL hygromycin B (ThermoFisher). Virus lytic replication in TREx-BCBL1-RTA cells was induced via addition of 2 µg/ml doxycycline hyclate (Sigma-Aldrich)^30^. HEK-293T cells were purchased from the ATCC and cultured in DMEM (Lonza) and supplemented with 10% FBS and 1% P/S. All cell lines were tested negative for mycoplasma.

### Antibodies and plasmid constructs

Antibodies used are listed as follows: EMG1 (Proteintech Cat#11965-1-AP, 1:500 WB), Lamin B1 (Sigma-Aldrich Cat # ZRB1143, 1:5000 WB), NOC4L (Proteintech Cat #17025-1-AP, WB 1:500), NOP14 ( Cat #26854-1-AP, WB 1:1000). FLAG (Sigma-Aldrich Cat#74425, 1:5000 WB), es19 (Proteintech Europe Cat#15085-1-AP, 1:500 WB), GAPDH (Proteintech Europe Cat#60004-1-Ig, 1:5000 WB), ORF57 (Santa Cruz Cat#sc-135747, 1:1000 WB), CDK1 (Abcam Cat# ab18, 1:5000 WB), ORF65 (Cambridge Research Biochemcials Cat#crb2005224, 1:100 WB), LANA (Sigma-Aldrich Cat#MABE1109, 1:50 IF).

Primers used for RT-qPCR are listen as follows: EMG1 (fwd: CCACCAGAGTTTGCTGATGC rvs: ATTCGGGTCTGGGGATT), GAPDH (fwd: TGTGGTCATGAGTCCTTCCACGAT; rvs: AGGGTCATC ATCTCTGCCCCCTC), ORF57 (fwd: GCCATAATCAAGCGTACTGG; rvs: GCAGACAAATATTGCGGTGT), ORF65 (fwd: AAGGTGAGAGACCCCGTGAT; rvs: TCCAGGGTATTCATGCGAGC), UBE2D2 (fwd: GTACTCTTGTCCATCTGTTCTCTG; rvs: CCATTCCCGAGCTATTCTGTT), Ywhaz (fwd: AAGACAGCACGCT AATAATGC; rvs: TTGGAAGGCCGGTTAATTTTC).

The packaging plasmids pVSV.G and psPAX2 were a gift from Dr Edwin Chen (University of Westminster, London). EMG1 shRNA constructs were purchased from GE healthcare (KD1 clone ID: TRCN0000134434, KD2 clone ID: V3SVHSHC_8318171). EMG1 mutant and EMG1 overexpression constructs were synthesised by GENEWIZ (Azenta).

### FLAG affinity purification assay

TREx-BCBL Rta cell lines overexpressing FLAG-ORF11-2xStrep tagged PNO1 bait protein^19^ or a control TREx BCBL1-Rta cell line were seeded at 0.75 x 10^6^ cells/ml in 20 mL RPMI selection or growth media. Half the flasks were induced with doxycycline, whilst half remained latent. After 24 hours cell pellets were resuspended in 1 mL pre-cooled ribosomal lysis buffer (10 mM Tris (pH 7.6), 100 mM KCl, 2 mM MgCl_2_, 0.5% (v/v) NP-40, 1 mM DTT, 1x protease inhibitor, 1% (v/v) phosphatase inhibitor), kept on ice for 20 minutes then centrifuged at 5000 x g for 12 minutes at 4 °C. Fab-TRAP leads (Proteintech) (30 µL per sample) were washed with 1 mL lysis buffer (without protease inhibitor, phosphatase inhibitor and DTT) three times and combined with the lysate for 1 hour at 4 °C with rotation. Samples were washed three times with pre-cooled ribosomal wash buffer 1 (10mM Tris-HCl (pH 7.6), 100 mM KCl, 2 mM MgCl_2_) and once with pre-cooled ribosomal wash buffer 2 (10 mM Tris-HCl (pH 7.6), 2 mM MgCl_2_). The FLAG-antibody-antigen complexes were resuspended in 2x Laemmeli buffer, and the samples were eluted by heating to 95°C for 5 minutes to be analysed by SDS-PAGE gel and Western blotting.

### Immunoblotting

Cell lysates were separated using 8-12% polyacrylamide gels and transferred to Amersham Nitrocellulose Membranes (GE healthcare) via Trans-blot Turbo Transfer system (Bio-Rad). Membranes were blocked in TBS + 0.1% tween with 5% wt/vol dried skimmed milk powder. Membranes were probed with appropriate primary antibodies and secondary horseradish peroxidase conjugated IgG antibodies at 1/5000 (Dako Agilent). Proteins were detected with ECL Western Blotting Substrate (Promega) or SuperSignal™ West Femto Maximum Sensitivity Substrate (ThermoFisher) before visualisation with G box (Syngene).

### RNA extraction and qPCR

Total RNA was extracted using Monarch Total RNA Miniprep Kit (NEB) as per manufacturer’s protocol. 1 µg RNA was reverse transcribed using LunaScript RT SuperMix Kit (NEB). qPCR was performed using synthesised cDNA, GoTaq qPCR MasterMix (promega) and the appropriate primer. qPCR was performed on Rotorgene Q and analysed by the ΔΔCT method against a housekeeping gene as previously described before plotting as fold^31^.

### Lentiviral expression and shRNA constructs

Lentiviruses were generated by transfection of HEK-293T cells using two packaging plasmids pPAX.2 and pVSV-G, a gift from Dr Edwin Chen (University of Westminster). Per 12-well, 4 µl of lipofectamine 2000 (Thermo Scientific) were used together with 1.2 µg of pLKO.1 plasmid expressing shRNA against the protein of interest, 0.65 µg of pVSV.G, and 0.65 µg psPAX2. Eight hours post-transfection, media was changed with 1.5 mL of DMEM supplemented with 10% (v/v) FCS. Two days post-transfection, viral supernatants were harvested, filtered through a 0.45 µm filter (Merck Millipore) and immediately used for transductions of TREx BCBL1-RTA cells. Cells (500,000) in 12 well plates were infected by spin inoculation for 60 min at 800 x g at room temperature, in the presence of 8 μg/mL of polybrene (Merck Millipore). 3 µg/ml Puromycin (Gibco) was added 48 hours after transduction before knockdown analysed >8 days post transduction via qPCR and western blot if appropriate^32^.

### Global translation assay

TREx BCBL1-Rta cells stably expressing scrambled shRNA or shRNAs targeting EMG1 were seeded into a 6-well plate at 1 × 10^6^ cells/well with 2 mL RPMI selection media without methionine which was chased out over 1 hr. Cells were treated with Click-iT® L-azidohomoalanine (AHA, 40 µM, Thermo Fisher) for 3 hrs, then washed in 1 XPBS and fixed in PBS containing 4% (v/v) paraformaldehyde for 15 min. Cells were again washed in 1X PBS and permeabilised using 1x PBS containing 0.25% Triton X-100 for 15 min. AHA was stained with a Click-iT® reaction kit (Thermo Fisher) using an alkyne Alexa Fluor™ 488, 1 µM, as described by the manufacturer. Finally, cells were washed in PBS containing 1% BSA and resuspended in PBS containing 0.5% BSA before fluorescence quantification by CytoFlex S Benchtop Flow Cytometer (Beckman Coulter).

### Viral reinfection assays

TREx BCBL1-Rta cells stably expressing scrambled shRNA or shRNAs targeting EMG1 were seeded in 2 mL of fresh RPMI media (300,000 cells/ mL) without puromycin into a 6-well plate. Lytic replication was induced by adding doxycycline (2 µg/mL) for 72 hours, with one scr well left uninduced as a control. Cell pellets were centrifuged at 500 *x g* for 5 minutes at 4⁰C. The supernatant, containing any released virion particles, was added in a 1:1 ratio to 293T cells seeded in the wells of 6-well plates at 50% confluency. The cells were incubated for 48 hrs, then harvested and RNA was isolated from the cell pellets. Purified RNA was reverse transcribed, and samples were analysed by qPCR for the levels of ORF57 to determine infectious virion production^33^.

### Immunofluorescence

Viral supernatant produced from viral reinfection assays were added to HEK293T cells grown on sterilised glass coverslips and incubated for 48 hours. The cells were fixed in PBS containing 4% paraformaldehyde (v/v) for 15 minutes, then permeabilised in PBS containing 1% Triton X-100 for 15 minutes. All the following steps were incubated for 1 hour at 37⁰C. Coverslips were blocked in PBS containing 1% BSA, incubated in anti-LANA (1/50), then incubated in Alexa Fluor 546 (1/500) before being mounted onto slides using Vectashield hardset mounting medium containing DAPI. Slides were visualised on a Zeiss LSM880 laser scanning confocal microscope and processed using ZEN 2009 imaging software.

### Polysome profiling

TREx BCBL1-Rta cells expressing scr shRNA or EMG1 specific shRNAs were treated with cycloheximide (Sigma) at 100 μg/ml for 3 min. A total of ∼50 × 10^6^ cells were pelleted and washed (1 x PBS, 100 μg/ml cycloheximide) then lysed in ice cold buffer [50 mM Tris-HCl pH8, 150 mM NaCl, 10 mM MgCl_2_, 1 mM DTT, 1% IGEPAL, 100 μg/ml cycloheximide, Turbo DNase 24 U/mL (Invitrogen), RNasin Plus RNase Inhibitor 90 U (Promega), 1 × protease inhibitor cocktail (Roche)] for 45 min. Lysates were clarified by centrifugation (12,000 × g, 10 min, 4 °C) and the resulting supernatants applied to 5–45% continuous sucrose gradients (10 mM MgCl_2_, 50 mM Tris-HCl (pH 7.6), 150 mM NaCl, 1 mM DTT, 100 μg/ml cycloheximide and 1× protease inhibitor cocktail). Gradients were then subjected to ultracentrifugation (160,000 *× g,* 3 hrs, 4 °C) in SW-40 rotor. Fractions from each gradient were collected using a Gradient Fractionator (BioComp) and the RNA profile was measured by absorbance (254 nm) across the gradient in real time using an EM-1 Econo UV Monitor (Bio-Rad). All fractions with large polysomes, six or more ribosomes per mRNA, were pooled and precipitated [0.3M NaCl, 50% isopropanol] overnight at −80°C. A Trizol RNA extraction was carried out as per the manufacturer’s protocol and samples were analysed via RT-qPCR.

### Luciferase assays

Luciferase activity was detected using the Dual-Luciferase Reporter Assay System (Promega). HEK-293T cell lines stably expressing a scr shRNA, EMG1 shRNA plasmid, an EMG1 mutant or wild type EMG1 overexpression construct were seeded in triplicate in 12-well culture plates (Thermo Fisher) at a density of 3 × 10^5^ cells per well and grown for 24 hours. Cells were transfected with 100 ng of respective plasmids (diluted in 100 µL Opti-mem) using 3 µL Lipofectamine® 2000 (Invitrogen^TM^; diluted in 100 µL Opti-mem) and incubated for a further 24hrs. Media was removed from the wells and cells washed gently with 100 μl PBS. Aliquots (100 μL) of 1x passive lysis buffer were added to the cell monolayers and rocked for 20 min at room temperature and then 10 μL of each lysate was transferred to tissue culture treated white microplates (Greiner Bio-One). Luciferase measurements were carried out using a FLUOstar Optima microplate reader (BMG Labtech Ltd), with injectors 1 and 2 being used to dispense 50 μL of Luciferase Assay Reagent II and Stop & Glo Reagent respectively. Renilla luciferase activity was normalized to Firefly luciferase activity.

### Statistical analysis

Except otherwise stated, graphical data shown represent mean ± standard deviation of mean (SD) using three biologically independent experiments. Differences between means was analysed by unpaired Student’s t test calculated using Graphpad Prism 9 calculator. Statistics was considered significant at p < 0.05, with *p<0.05, **p<0.01, ***p0.001.

## References

1. Frank, J. (2000). The ribosome--a macromolecular machine par excellence. Chem Biol 7, R133–141.

2. Hinkson, I.V., and Elias, J.E. (2011). The dynamic state of protein turnover: It’s about time. Trends Cell Biol 21, 293–303. 10.1016/j.tcb.2011.02.002.

3. Genuth, N.R., and Barna, M. (2018). The Discovery of Ribosome Heterogeneity and Its Implications for Gene Regulation and Organismal Life. Mol Cell 71, 364–374. 10.1016/j.molcel.2018.07.018.

4. Xue, S., and Barna, M. (2012). Specialized ribosomes: a new frontier in gene regulation and organismal biology. Nat Rev Mol Cell Biol 13, 355–369. 10.1038/nrm3359.

5. Shi, Z., Fujii, K., Kovary, K.M., Genuth, N.R., Rost, H.L., Teruel, M.N., and Barna, M. (2017). Heterogeneous Ribosomes Preferentially Translate Distinct Subpools of mRNAs Genome-wide. Mol Cell 67, 71–83 e77. 10.1016/j.molcel.2017.05.021.

6. Hopes, T., Norris, K., Agapiou, M., McCarthy, C.G.P., Lewis, P.A., O’Connell, M.J., Fontana, J., and Aspden, J.L. (2022). Ribosome heterogeneity in Drosophila melanogaster gonads through paralog-switching. Nucleic Acids Res 50, 2240–2257. 10.1093/nar/gkab606.

7. Gupta, V., and Warner, J.R. (2014). Ribosome-omics of the human ribosome. RNA 20, 1004–1013. 10.1261/rna.043653.113.

8. Simsek, D., Tiu, G.C., Flynn, R.A., Byeon, G.W., Leppek, K., Xu, A.F., Chang, H.Y., and Barna, M. (2017). The Mammalian Ribo-interactome Reveals Ribosome Functional Diversity and Heterogeneity. Cell 169, 1051–1065 e1018. 10.1016/j.cell.2017.05.022.

9. Simsek, D., and Barna, M. (2017). An emerging role for the ribosome as a nexus for post-translational modifications. Curr Opin Cell Biol 45, 92–101. 10.1016/j.ceb.2017.02.010.

10. Roundtree, I.A., Evans, M.E., Pan, T., and He, C. (2017). Dynamic RNA Modifications in Gene Expression Regulation. Cell 169, 1187–1200. 10.1016/j.cell.2017.05.045.

11. Kampen, K.R., Sulima, S.O., Vereecke, S., and De Keersmaecker, K. (2020). Hallmarks of ribosomopathies. Nucleic Acids Res 48, 1013–1028. 10.1093/nar/gkz637.

12. Stern-Ginossar, N., Thompson, S.R., Mathews, M.B., and Mohr, I. (2019). Translational Control in Virus-Infected Cells. Cold Spring Harb Perspect Biol 11. 10.1101/cshperspect.a033001.

13. Walsh, D., Mathews, M.B., and Mohr, I. (2013). Tinkering with translation: protein synthesis in virus-infected cells. Cold Spring Harb Perspect Biol 5, a012351. 10.1101/cshperspect.a012351.

14. Ganem, D. (2010). KSHV and the pathogenesis of Kaposi sarcoma: listening to human biology and medicine. Journal of Clinical Investigation 120, 939–949. 40567 [pii]10.1172/JCI40567.

15. Dittmer, D.P., and Damania, B. (2019). Kaposi’s Sarcoma-Associated Herpesvirus (KSHV)- Associated Disease in the AIDS Patient: An Update. Cancer Treat Res 177, 63–80. 10.1007/978-3-030-03502-0_3.

16. Broussard, G., and Damania, B. (2020). Regulation of KSHV Latency and Lytic Reactivation. Viruses 12. 10.3390/v12091034.

17. Giffin, L., and Damania, B. (2014). KSHV: pathways to tumorigenesis and persistent infection. Adv Virus Res 88, 111–159. 10.1016/B978-0-12-800098-4.00002-7.

18. Arias, C., Weisburd, B., Stern-Ginossar, N., Mercier, A., Madrid, A.S., Bellare, P., Holdorf, M., Weissman, J.S., and Ganem, D. (2014). KSHV 2.0: a comprehensive annotation of the Kaposi’s sarcoma-associated herpesvirus genome using next-generation sequencing reveals novel genomic and functional features. PLoS Pathog 10, e1003847. 10.1371/journal.ppat.1003847.

19. Murphy, J.C., Harrington, E.M., Schumann, S., Vasconcelos, E.J.R., Mottram, T.J., Harper, K.L., Aspden, J.L., and Whitehouse, A. (2023). Kaposi’s sarcoma-associated herpesvirus induces specialised ribosomes to efficiently translate viral lytic mRNAs. Nat Commun 14, 300. 10.1038/s41467-023-35914-5.

20. Haag, S., Kretschmer, J., and Bohnsack, M.T. (2015). WBSCR22/Merm1 is required for late nuclear pre-ribosomal RNA processing and mediates N7-methylation of G1639 in human 18S rRNA. RNA 21, 180–187. 10.1261/rna.047910.114.

21. Wurm, J.P., Meyer, B., Bahr, U., Held, M., Frolow, O., Kotter, P., Engels, J.W., Heckel, A., Karas, M., Entian, K.D., and Wohnert, J. (2010). The ribosome assembly factor Nep1 responsible for Bowen-Conradi syndrome is a pseudouridine-N1-specific methyltransferase. Nucleic Acids Res 38, 2387–2398. 10.1093/nar/gkp1189.

22. Meyer, B., Wurm, J.P., Kotter, P., Leisegang, M.S., Schilling, V., Buchhaupt, M., Held, M., Bahr, U., Karas, M., Heckel, A., et al. (2011). The Bowen-Conradi syndrome protein Nep1 (Emg1) has a dual role in eukaryotic ribosome biogenesis, as an essential assembly factor and in the methylation of Psi1191 in yeast 18S rRNA. Nucleic Acids Res 39, 1526–1537. 10.1093/nar/gkq931.

23. Cheng, J., Lau, B., Thoms, M., Ameismeier, M., Berninghausen, O., Hurt, E., and Beckmann, R. (2022). The nucleoplasmic phase of pre-40S formation prior to nuclear export. Nucleic Acids Res 50, 11924–11937. 10.1093/nar/gkac961.

24. Leulliot, N., Bohnsack, M.T., Graille, M., Tollervey, D., and Van Tilbeurgh, H. (2008). The yeast ribosome synthesis factor Emg1 is a novel member of the superfamily of alpha/beta knot fold methyltransferases. Nucleic Acids Res 36, 629–639. 10.1093/nar/gkm1074.

25. Armistead, J., Khatkar, S., Meyer, B., Mark, B.L., Patel, N., Coghlan, G., Lamont, R.E., Liu, S., Wiechert, J., Cattini, P.A., et al. (2009). Mutation of a gene essential for ribosome biogenesis, EMG1, causes Bowen-Conradi syndrome. Am J Hum Genet 84, 728–739. 10.1016/j.ajhg.2009.04.017.

26. Lowry, R.B., Innes, A.M., Bernier, F.P., McLeod, D.R., Greenberg, C.R., Chudley, A.E., Chodirker, B., Marles, S.L., Crumley, M.J., Loredo-Osti, J.C., et al. (2003). Bowen-Conradi syndrome: a clinical and genetic study. Am J Med Genet A 120A, 423–428. 10.1002/ajmg.a.20059.

27. Warda, A.S., Freytag, B., Haag, S., Sloan, K.E., Gorlich, D., and Bohnsack, M.T. (2016). Effects of the Bowen-Conradi syndrome mutation in EMG1 on its nuclear import, stability and nucleolar recruitment. Hum Mol Genet 25, 5353–5364. 10.1093/hmg/ddw351.

28. Zhao, Y., Rai, J., Yu, H., and Li, H. (2022). CryoEM structures of pseudouridine-free ribosome suggest impacts of chemical modifications on ribosome conformations. Structure 30, 983–992 e985. 10.1016/j.str.2022.04.002.

29. Babaian, A., Rothe, K., Girodat, D., Minia, I., Djondovic, S., Milek, M., Spencer Miko, S.E., Wieden, H.J., Landthaler, M., Morin, G.B., and Mager, D.L. (2020). Loss of m(1)acp(3)Psi Ribosomal RNA Modification Is a Major Feature of Cancer. Cell Rep 31, 107611. 10.1016/j.celrep.2020.107611.

30. Schumann, S., Jackson, B.R., Yule, I., Whitehead, S.K., Revill, C., Foster, R., and Whitehouse, A. (2016). Targeting the ATP-dependent formation of herpesvirus ribonucleoprotein particle assembly as an antiviral approach. Nat Microbiol 2, 16201. 10.1038/nmicrobiol.2016.201.

31. Harper, K.L., Mottram, T.J., Anene, C.A., Foster, B., Patterson, M.R., McDonnell, E., Macdonald, A., Westhead, D., and Whitehouse, A. (2022). Dysregulation of the miR-30c/DLL4 axis by circHIPK3 is essential for KSHV lytic replication. EMBO Rep, e54117. 10.15252/embr.202154117.

32. Baquero-Perez, B., Antanaviciute, A., Yonchev, I.D., Carr, I.M., Wilson, S.A., and Whitehouse, A. (2019). The Tudor SND1 protein is an m(6)A RNA reader essential for replication of Kaposi’s sarcoma-associated herpesvirus. Elife 8. 10.7554/eLife.47261.

33. Manners, O., Baquero-Perez, B., Mottram, T.J., Yonchev, I.D., Trevelyan, C.J., Harper, K.L., Menezes, S., Patterson, M.R., Macdonald, A., Wilson, S.A., et al. (2023). m(6)A Regulates the Stability of Cellular Transcripts Required for Efficient KSHV Lytic Replication. Viruses 15. 10.3390/v15061381.

